# Spike-in normalization for single-cell RNA-seq reveals dynamic global transcriptional activity mediating anti-cancer drug response

**DOI:** 10.1101/2021.01.24.427929

**Authors:** Xin Wang, Jane Frederick, Hongbin Wang, Sheng Hui, Vadim Backman, Zhe Ji

## Abstract

The transcriptional plasticity of cancer cells promotes intercellular heterogeneity in response to anti-cancer drugs and facilitates the generation of subpopulation surviving cells. Characterizing single-cell transcriptional heterogeneity after drug treatments can provide mechanistic insights into drug efficacy. Here we used single-cell RNA-seq to examine transcriptomic profiles of cancer cells treated with paclitaxel, celecoxib, and the combination of the two drugs. By normalizing the expression of endogenous genes to spike-in molecules, we found that celluar mRNA abundance shows dynamic regulation after drug treatment. Using a random forest model, we identified gene signatures classifying single cells into three states: transcriptional repression, amplification, and control-like. Treatment with paclitaxel or celecoxib alone generally repressed gene transcription across single cells. Interestingly, the drug combination resulted in transcriptional amplification and hyperactivation of mitochondrial oxidative phosphorylation pathway linking to enhanced cell killing efficiency. Finally, we identified a regulatory module enriched with metabolism and inflammation-related genes activated in a subpopulation of paclitaxel-treated cells, the expression of which predicted paclitaxel efficacy across cancer cell lines and *in vivo* patient samples. Our study highlights the dynamic global transcriptional activity driving single-cell heterogeneity during drug response and emphasizes the importance of adding spike-in molecules to study gene expression regulation using single-cell RNA-seq.

## INTRODUCTION

A major challenge of cancer therapy is the acquired resistance of cancer cells to chemotherapy drugs and associated disease relapse. A driver of drug resistance is intercellular heterogeneity of gene expression resulting from genetic mutations and epigenetic alterations (1,2). The preexisting and rewired gene expression programs of cancer cells determine their ultimate fates after drug treatment. Emerging evidence shows that even genetically identical cells show transcriptional heterogeneity in response to external stimuli under both physiological and *in vitro* conditions, because of their transcriptional plasticity (3,4). Characterizing single-cell transcriptomic dynamics after drug treatment can provide mechanistic insights into cell fate decisions and identify predictive biomarkers of efficacy (5).

Paclitaxel is a commonly used chemotherapy drug for treating diverse human cancers, such as ovarian, breast, and lung cancers. It binds to the microtubule polymer and disrupts its disassembly, which triggers mitotic arrest and apoptosis (6). However, most patients develop chemoresistance after several sessions of treatment. Many studies have been devoted to identifying the molecular mechanisms mediating paclitaxel resistance and developing combinatory therapeutic strategies (7,8). For example, synergistic inhibition of NF-κB and PI3K signaling pathways can sensitize paclitaxel-resistant cancer cells (9,10). Combination treatment with the nonsteroidal anti-inflammatory drug celecoxib (a COX-2 inhibitor) can increase paclitaxel’s efficacy at killing cancer cells (11–13).

Despite extensive effort, the molecular mechanisms regulating paclitaxel efficacy remain elusive. A limitation is that many studies were carried out using bulk cancer cells, without considering the contribution of intercellular heterogeneity. Recently, using partial wave spectroscopic (PWS) microscopy, we observed that treatment with paclitaxel increases the packing density scaling of chromatin domains and their intercellular heterogeneity of surviving subpopulations of cancer cells, suggesting that surviving cells exhibit a phenotype consistent with enhanced single-cell transcriptional heterogeneity (14). On the other hand, celecoxib treatment decreases the chromatin packing density scaling within chromatin domains of cells. Due to the opposite effects of paclitaxel and celecoxib on chromatin packing in the nucleus, we hypothesized that their combination may induce a *de novo* transcriptional program promoting cancer cell death.

To obtain molecular insights into the drug response, we performed full-length single-cell RNA sequencing (scRNA-seq) with Smart-seq2 (15) and profiled the transcriptomes of several hundred single cells at different time points after treatment with paclitaxel, celecoxib, and the combination of the two drugs, respectively. To compare the global transcriptomic levels across single cells, we added a constant amount of polyA-tailed spike-in molecules to each cell during the library preparation (16). By normalizing endogenous gene expression levels to those of spike-in molecules, we found that global transcriptomic levels are heterogeneous across single cells and are dynamically regulated after drug treatment.

Paclitaxel treatment alone generally repressed transcriptomic levels, while its combination with celecoxib resulted in transcriptional amplification. We developed a random forest model and classified single cells based on their transcriptional states. The model revealed that the downregulation of the cell cycle pathway is associated with transcriptional repression, and the hyperactivation of mitochondrial oxidative phosphorylation (OXPHOS) contributes to the transcriptional amplification and enhanced cell killing efficacy after the drug combination. Furthermore, we identified a coherent gene module regulating cellular metabolism and inflammation, the higher expression of which predicts worse paclitaxel response in cancer cell lines and patients. Altogether, our study highlights the unprecedented effects of single-cell heterogeneity on global transcriptional activity in regulating the anti-cancer drug response.

## MATERIALS AND METHODS

### Cell culture

A2780 ovarian endometroid adenocarcinoma cells were a gift from Dr. Chia-Peng Huang Yang at the Albert Einstein College of Medicine obtained from Dr. Elizabeth de Vries at the University Medical Center Groningen. The cells were cultured in RPMI 1640 medium (Thermo Fisher Scientific, Waltham, MA) supplemented with 10% fetal bovine serum (Thermo Fisher Scientific, Waltham, MA) on 35-mm six-well glass bottom plates (Cellvis, Mountain View, CA) until 60 to 85% confluent. All cells were given at least 24 hours to readhere before drug treatment.

### Cell growth and apoptosis experiments

Cells were treated with 75 μM celecoxib, 5 nM paclitaxel, or a combination of 75 μM celecoxib and 5 nM paclitaxel for 48 hours. Cells were then imaged to determine the percent coverage of the well for each treatment. To determine the amount of cell growth inhibition based on a treatment, the amount of cell coverage was normalized to the control group. Apoptosis staining was performed using the CellEvent Caspase-3/7 Green Flow Cytometry Assay Kit (Thermo Fisher Scientific, Waltham, MA). Stained cell suspensions were measured with the BD LSRFortessa Cell Analyzer (BD Biosciences, San Jose, CA) at the Northwestern University Flow Cytometry Core Facility.

### scRNA-seq Library preparation using SMART-seq2

Cells were treated with 75 μM celecoxib for 2 and 16 hours, 5 nM paclitaxel for 16 and 48 hours, or a combination of 75 μM celecoxib and 5 nM paclitaxel for 16 hours prior to trypsinization and resuspension in growth medium. Cell suspensions were sorted with a C1 single-cell capture system (Fluidigm, South San Francisco, CA) by the University of Illinois at Chicago Genomics Core. scRNA-seq libraries were prepared according to the Smart-seq2 protocol and sequenced using the NextSeq 500 Sequencing System (Illumina, San Diego, CA) by the University of Illinois at Chicago Sequencing Core. A predesigned set of three polyA-tailed spike-in RNAs (sequences are shown in Table S1) was added to each well during the library preparation.

### scRNA-seq data processing

We trimmed the adapter sequences of raw sequencing reads using the Trim Galore software (https://www.bioinformatics.babraham.ac.uk/projects/trim_galore/). We used RSEM (17) to calculate RNA expression levels using scRNA-seq data. We created an RSEM index with the combination of the human hg38 transcriptome (GENCODE version 28 (18)) and 3 spike-in molecules. We calculated RNA expression using RSEM 1.3.0 with the default parameters and Bowtie2 (19) as the aligner. We normalized the expression levels of endogenous genes to that of the highest expressed spike-in molecule. Given *M* single cells, to control the dynamic regulation of endogenous gene expression in single cell *k*, the TPM value of gene *j* is normalized to that of spike-in 1 (*s*), which is the highest spike-in molecule. The normarlized expression of gene *j* in the cell *k* (*N_jk_*) was calculated as:

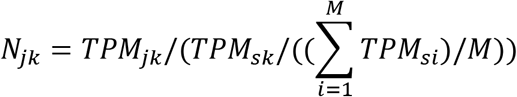

As the RNA capturing complexity of the scRNA-seq library is ~10% of bulk RNA-seq (20), we divided the above-normalized TPM values by 10, and the expression level of gene *j* in cell *k* was quantified as the E-value *E_jk_*:

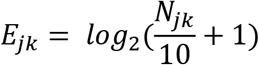

We performed quality-control of the scRNA-seq data and filtered out cells with housekeeping genes poorly detected (averaged E-value of the genes <1) (21). The housekeeping genes were defined in (22). We only included genes detected in >30% of single cells in at least one experimental condition in further analyses.

### Principal component analysis and random forest modeling to classify the cell states

We used the averaged E-values of a gene across single cells to indicate its expression upon drug treatment. We selected genes showing significant differential expression after drug treatment (>1.5-fold change compared to control cells) for the principal component analysis and building the random forest model. For each gene, the control-normalized E-values were used as the input for the analyses.

To build the random forest model, we used the control cells, cells treated with celecoxib for 16 h, and cells treated with both paclitaxel and celecoxib to represent the control-like, transcriptional repression and activation states, respectively. We randomly selected 2/3 of single cells from these treatment conditions to train the model with 5-fold cross-validation. The classification of the remaining 1/3 of the cells was used to measure the algorithm performance using ROC curves. Genes with an MDA value >0 from the model contributed significantly to the classification. Finally, the model was used to classify all single cells into all three states with 3 corresponding *P*-values. The sum of the 3 *P*-values was 1, and the state with the largest *P*-value was defined as the classified state.

### Differential gene expression and pathway analysis

We grouped genes into 3 coherent clusters based on their relative differential expression changes after drug treatment. We used the average E-values across single cells in a drug treatment condition to indicate the expression level. Genes in cluster I were down-regulated >1.5-fold at 16 h of celecoxib treatment and up-regulated >1.5-fold at 16 h of combination drug treatment. Genes in cluster II were down-regulated at 16 h of celecoxib treatment and showed no significant expression change after combination treatment. Genes in cluster III were up-regulated >1.5-fold after combination treatment and showed no significant regulation after celecoxib treatment. The gene ontology analyses were performed using the Database for Annotation, Visualization, and Integrated Discovery (DAVID) (23).

For a geneset regulating a biological process, we compared its expression levels across single cells after each drug treatment. Taking the OXPHOS pathway for example, we used the expressed genes in GO:0006119 for the analyses. Suppose there are *G* genes in the geneset. For each gene *j*, we first normalized its E-value in cell *k* to the mean of the control cells as *M_jk_*. The relative expression of the geneset in cell *k* was then calculated as the mean(*M_jk_*) _*j=1, 2… G*_.

Using this method, we calculated the relative expression of genesets encoding complex I (GO:0005747), complex II (GO:0005749), complex III (GO:0005750), complex IV (GO:0005751), and ATP synthase of the OXPHOS pathway. We used genes in GO:0007049 for the analyses of cell cycle regulation. For genes showing unique activation in a particular cell cycle phase, we used previously defined gene lists (24). To compare the relative expression of a set of metabolism genes, we used the following paclitaxel-activated genes in the GO:0055114: oxidation-reduction process geneset for analysis, including TM7SF2, PHYHD1, ACADSB, HMGCR, HSD17B14, ALDH1A1, MTHFD2, TP53I3, PLOD1, NOS3, CCS, GPX8, HSD17B8, PTGR1, PTGR2, DHRS12, SCD, FADS1, DECR2, PHYH, VAT1, MSRB2, COQ6, BLVRA, SQLE, ALDH2, PHGDH, ACAD11.

### Calculation of the paclitaxel response index

We downloaded the RNA expression data measured by Affymetrix microarray and the corresponding annotations of 1,037 cancer cell lines from the CCLE (25). Based on the averaged E-values across single cells, 177 genes were up-regulated (>1.5-fold) in the control-like cell subpopulation after 48 h of paclitaxel treatment. As these genes are co-activated after paclitaxel treatment, we next examined whether they form a coherent transcriptional module and their baseline expression levels are correlated across cancer cell lines. Using the CCLE data and an iterative computational approach, we identified 73 genes with a significant positive correlation of expression. The approach is as follows.

We analyzed the RNA expression levels of the 177 paclitaxel-activated genes using CCLE data. First, we required that a gene included for further analysis should be expressed in >50% of cancer cells and show variable expression levels across all 1,037 cancer cells. We used the MAS5 algorithm to determine whether a gene is expressed in a cell. The difference between the 90th percentile and the 10th percentile of the gene was required to be greater than 3-fold. This step filtered out genes showing lineage-specific or constitutive expression across cancer cells.

Second, for the geneset consisting of remaining genes from the above step, we calculated its relative expression across all 1,037 cancer cells using a similar method as described above for the pathway analyses. Given *G* genes in the geneset, for each gene *j*, we first normalized its log2-based expression level in cell *k* to the median of all cancer cells as *M_jk_*, and the relative expression of the geneset in cell *k* was calculated as the median(*M_jk_*)_*j=1, 2… G*_.

Third, we calculated the Spearman correlation coefficients between the geneset expression and the expression levels of each gene across all 1,037 cancer cells. We removed genes with a coefficient value <0.1 from the geneset. Using an iterative repetition of step 2 and step 3, we obtained 73 genes showing a significant positive correlation with each other. We then calculated the relative expression of the entire set of 73 genes as the paclitaxel response index across cancer cell lines. The 73 genes include SLC25A21, PTGR2, SCD, ALDH1A1, PLTP, ALDH2, FKBP1B, TP53I3, SERPING1, C6orf1, VAT1, PLOD1, IFI27L2, ARMCX3, BLVRA, SPA17, EFHC1, SBF2, RAB13, CDC42EP5, TNNC1, FBXO2, TLCD1, PLEKHA1, SAT1, SEMA3E, USP32, ABI2, PHYHD1, MYL5, DHRS12, OCEL1, HMGCL, CORO1B, GGPS1, ITM2B, NOS3, DECR2, ECI1, C9orf16, NEK3, SUCLG2, CRYL1, TST, ACER3, SERPINB6, CD46, YIPF3, SUMF2, SIL1, GRN, MSRB2, PHYH, TNFRSF10B, ETV4, DUSP6, S100A4, IFITM1, IFITM2, IFITM3, CA2, GJA1, IFI35, RAB32, LGALS1, HSPA1A, MYO1B, GPX8, ANXA1, PTGR1, RRAS, TNFRSF1A, S100A11.

### Cancer patient data analysis

We downloaded two clinical cohorts of breast cancer patient data from the NCBI Gene Expression Omnibus (GEO) database, including GSE25066 (Hatzis dataset (26)) and GSE32646 (Miyake dataset (27)). We used the R package affxparser to read and analyze RNA expression levels measured by the microarray data. Because the analysis is for breast cancer patients and RNA expression represents the averaged signals across cancer cells and stromal cells, we required that genes used to calculate the paclitaxel response index should be expressed in >50% of CCLE breast cancer cell lines. We used the following 51 genes to calculate the paclitaxel response index in breast cancer patients: S100A11, GRN, HSPA1A, PLOD1, ANXA1, LGALS1, IFITM2, ALDH2, IFITM1, RAB13, GGPS1, HMGCL, S100A4, PHYH, SAT1, BLVRA, RAB32, C9orf16, DHRS12, MYL5, SPA17, OCEL1, FKBP1B, ABI2, CD46, TNFRSF1A, VAT1, DUSP6, IFI35, TST, ECI1, TP53I3, NEK3, SERPINB6, USP32, IFITM3, MYO1B, SUCLG2, RRAS, YIPF3, ITM2B, ARMCX3, SIL1, MSRB2, PLEKHA1, FBXO2, DECR2, EFHC1, SLC25A21, CRYL1, CORO1B.

## RESULTS

### scRNA-seq of cancer cells at different time points after drug treatments

We treated A2780 ovarian cancer cells with paclitaxel, celecoxib, and their combination (Figure 1A). Consistent with previous reports, co-treatment of celecoxib increased paclitaxel-induced apoptosis (Figure 1B-C). To characterize intercellular transcriptional heterogeneity mediating the drug response, we performed full-length scRNA-seq with Smart-seq2 using the Fluidigm C1 system for a total of 372 live cells at different time points after drug treatment, including 16 h and 48 h with paclitaxel, 2 h and 16 h with celecoxib, and 16 h with the two drugs (Figures 1A and S1A). We obtained scRNA-seq data for ~60 single cells from each experimental condition. We used Smart-seq2 for the experiment because it has the highest sensitivity to capture expressed transcripts compared to other scRNA-seq techniques (28). We picked the early time points because initial gene expression changes after drug treatment are crucial for determining cell fates (29), and they are representative of the dynamic chromatin packing density revealed by our PWS experiments. To examine the regulation of global transcriptomic abundance in single cells and perform quality control of the scRNA-seq experiment, we used a pre-designed spike-in set consisting of three polyA-tailed RNAs with variable lengths and concentrations (Table S1). We added an equal volume of the spike-in set to each well of the 384-well plate during the library preparation using the Fluidigm C1 system.

**Figure 1.**
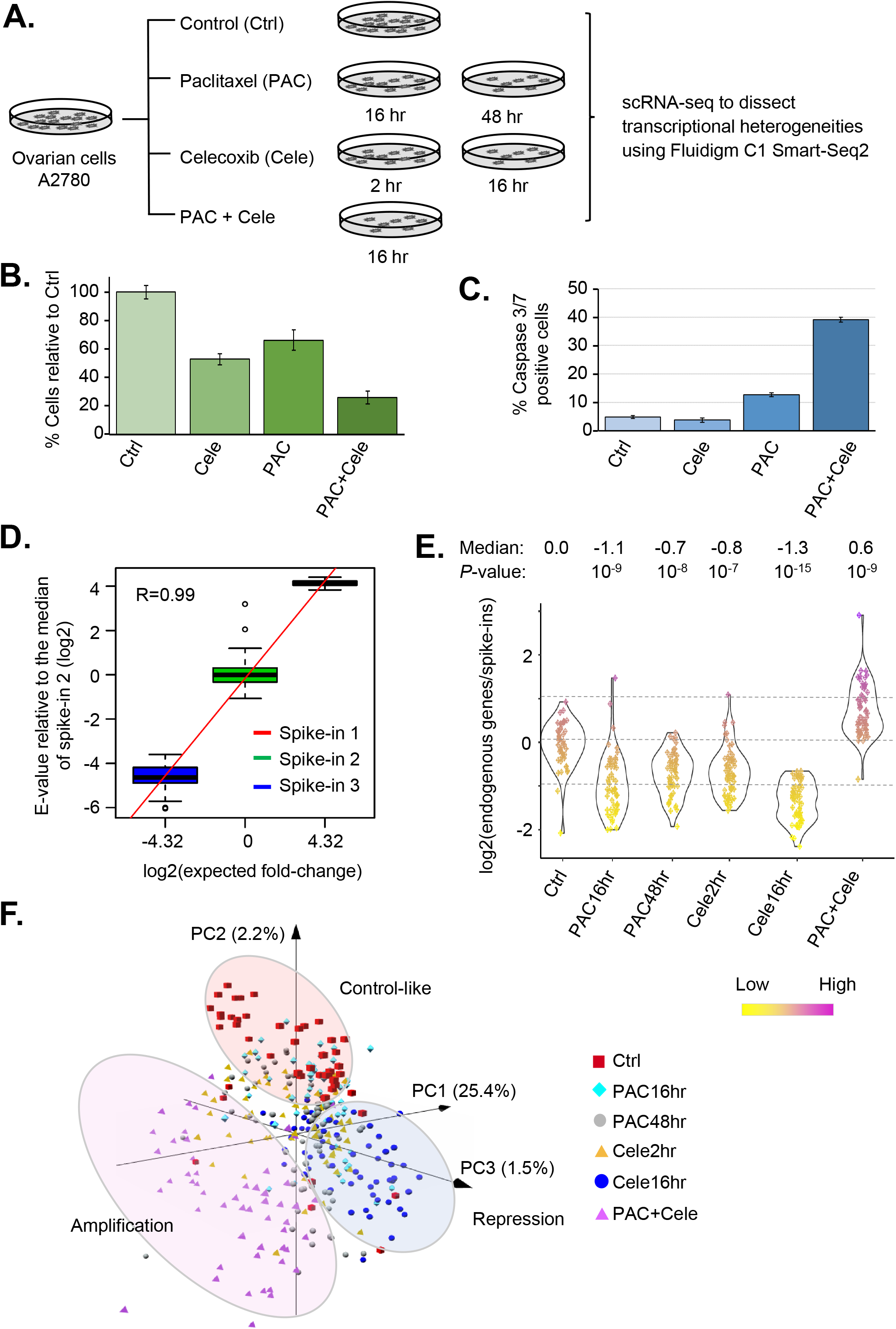
scRNA-seq to examine transcriptional heterogeneities after drug treatment. (A) We treated A2780 ovarian cancer cells with paclitaxel (PAC), celecoxib (Cele), and the combination of the two drugs, and collected the cells at different time points for scRNA-seq using Smart-Seq2 on the Fluidigm C1 system. (B) The relative cell growth after 48 h drug treatments as compared to controls. (C) The fraction of apoptotic cells (caspase 3/7+) after 48 h drug treatment. (D) The correlation between expected fold differences of spike-in molecules vs. those calculated by scRNA-seq data across single cells. The expression values were normalized to the median of spike-in 2. The Pearson correlation coefficient is indicated in the plot. (E) The relative transcriptomic abundance across single cells is measured as the ratio between the number of reads mapped to endogenous mRNAs and the number mapped to the spike-in molecules. The log2(ratio) values were normalized to the median of the control cells. Each dot in the plot represents a single cell. The Wilcoxon rank sum test *P*-values comparing drug treatment groups vs. the control are shown. (F) principal component analysis of single cells treated with drugs at different time points. We used expression profiles of genes showing significant differential expression in at least one drug treatment condition for this analysis. The percentage of variance explained by the principal component is shown in parentheses.

For each single-cell library, sequencing reads were mapped to the hybrid reference transcriptome (GENCODE-defined endogenous transcripts + spike-in RNA sequences) to quantify the RNA expression. The fold expression differences of spike-in RNAs are consistent with the pre-designed conditions (Figures 1D and S2), indicating that our scRNA-seq experiment quantitatively measures RNA expression levels. For each endogenous gene, we normalized its “transcript per million” (TPM) value to that of the highest expressed spike-in RNA in the same cell, and used the normalized value to indicate its expression level. We removed 30 poor quality cells with a low number of housekeeping genes detected (Table S2, Figure S1B and see Methods for details) (20,21). We retained 338 single cells for further analyses. In addition, we required that a gene included in the analyses should be detected in at least 30% of single cells from at least one experimental condition. After these quality-control steps, we were able to study the dynamic expression of ~9,600 genes.

### Dynamic regulation of global transcriptomic levels

To measure the relative transcriptomic amount in a single cell, we calculated the ratio between the number of reads mapped to endogenous mRNAs and the number mapped to spike-in RNAs. In each experimental condition, the overall mRNA abundance varied >3-fold across single cells indicating intercellular heterogeneity (Figure 1E) (30). If we consider the median value across single cells in a condition, the global transcriptomic level is repressed after paclitaxel or celecoxib treatment alone. As compared to the control cells, the overall mRNA amount decreases ~2.1-fold at 16 hr of paclitaxel treatment (*P* < 10^−9^), and reduces ~2.4-fold at 16 hr with celecoxib (*P* < 10^−15^) (Figure 1E). Interestingly, after 16 h co-treatment of paclitaxel and celecoxib, the transcriptomic amount was up-regulated ~1.5-fold (*P* < 10^−9^) (Figure 1E). These results suggest that the combination of the two drugs switches the transcriptional repression induced by a single drug to the transcriptional amplification state. The same result was produced when we used a different computational approach to measure relative transcriptomic levels, by calculating the ratio between the sum of TPM values of the top 5,000 expressed mRNAs vs. that of the highest spike-in RNA (Figure S3).

Next, using the principal component analysis, we clustered single cells based on their transcriptomic profiles. The unbiased clustering showed that the single cells are likely in three different states: the control-like state (e.g., untreated cancer cells), the transcriptional repression state (e.g., cells with 16 hr celecoxib treatment), and the transcriptional amplification state (e.g., cells co-treated with paclitaxel and celecoxib) (Figure 1F). Cells from other conditions, especially with paclitaxel treatment alone, consist of mixtures corresponding to different states (Figure 1F).

### A random forest model classifies single cells into transcriptional repression, amplification, and control-like states

To further characterize the intercellular heterogeneity, we developed a supervised learning model using the random forest method to classify single cells into the three states based on their transcriptomic profiles. We used gene expression profiles of cells treated with celecoxib for 16 h, cells co-treated with the two drugs, and untreated cancer cells to represent transcriptional repression, amplification, and control-like states, respectively, because the cells from these conditions tend to be homogenous (Figure 1F). We used expression profiles from a randomly selected 2/3 of cells in each group as the training set, and profiles of the remaining 1/3 as the testing set to evaluate the algorithm performance. The random forest model performed the feature gene selection and classified cell states with high accuracy (area under the receiver operating characteristic (ROC) curve (AUC) >0.95 for each of three states) (Figure 2A).

**Figure 2.**
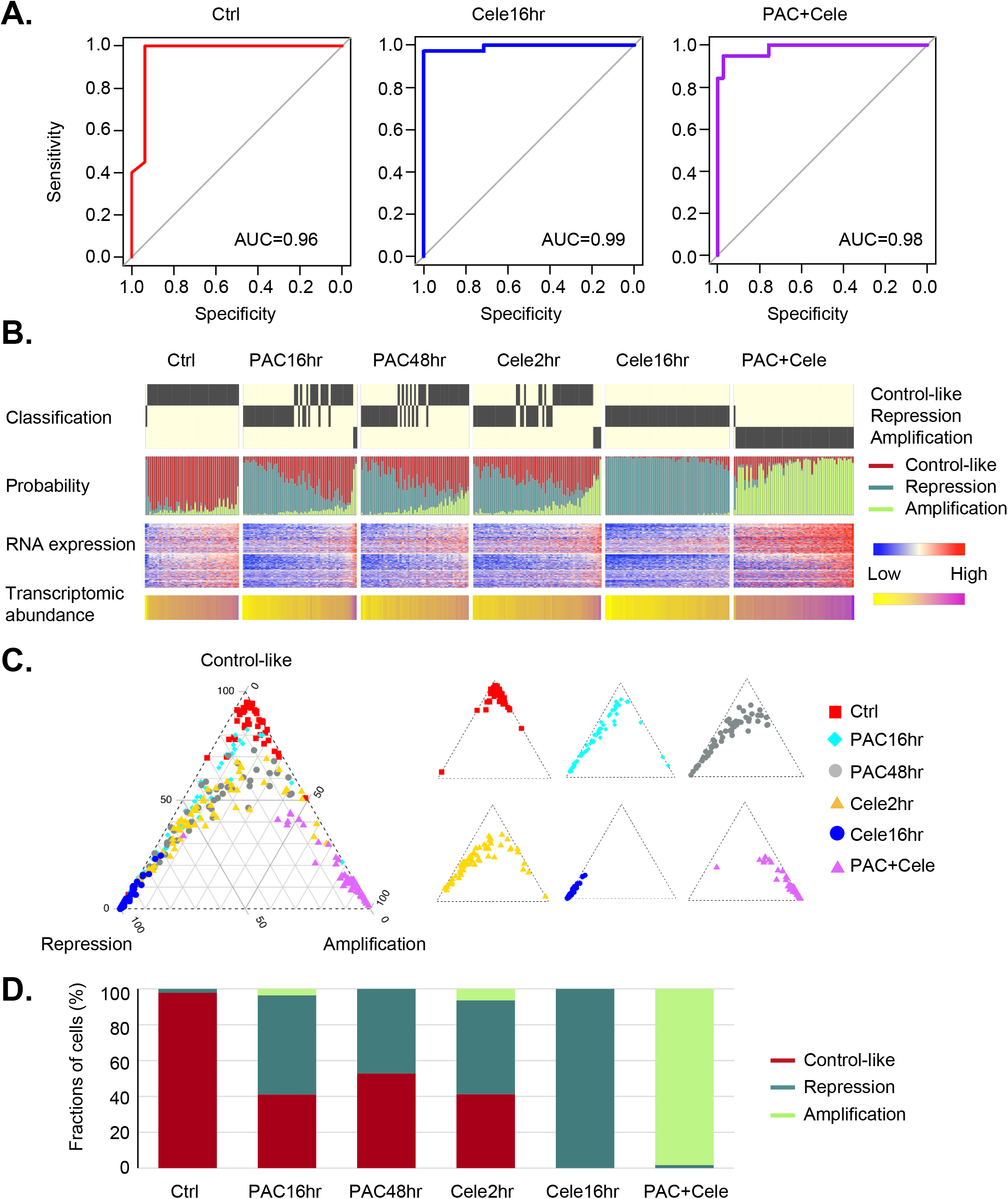
A random forest model classifies single cells into three different states: transcriptional repression, amplification, and control-like. (A) We used transcriptomic profiles of cells with 16 h of celecoxib, 16 h of combination treatment, and control cells, to represent the transcriptional repression, amplification, and control-like states, respectively. We used 2/3 of single cells for training and the remaining 1/3 of cells for testing. The algorithm performance when classifying the three states based on the testing set was measured using ROC curves, and the area under the ROC curves (AUC) is shown in each plot. (B) Classifying single cells into three states based on their gene expression profiles using the random forest model. For each single cell, the transcriptomic abundance (as in Figure 1E), RNA expression profile (normalized E-values by mean of control cells), predicted *P*-values of the three states, and the decision state are shown. Each column in the plot represents a single cell. (C) The triangle plot showing the predicted *P*-values of the three states across single cells. The pooled single cells are shown on the left, and cells with different treatment conditions are shown on the right. (D) Fraction of cells classified into the three states under different drug treatment conditions.

We then applied the random forest model and classified all single cells based on their RNA expression profiles (Figure 2B-C). This classification process refined the dynamic compositions of cells from different states after paclitaxel treatment. At 16 h with paclitaxel, 55% of cells show transcriptional repression, 4% show transcriptional amplification and 41% are control-like (Figure 2D). At 48 h, cells treated with paclitaxel remain in a mixture of states, including 47% in the transcriptional repression state and 53% in the control-like state. These results highlight the intercellular transcriptional heterogeneity in response to paclitaxel and are consistent with our PWS microscopy results showing changes in chromatin packing density (14).

### Regulation of OXPHOS and cell cycle genes is a major determinant of transcriptomic state

In addition to the global regulation of transcriptomic abundance, individual genes show variable expression changes after drug treatment. We partitioned drug-responsive genes into three co-regulatory clusters based on their relative expression across single cells (Figure 3A and Table S3). For each gene, we annotated its contribution to the random forest model indicated by the mean decrease in accuracy (MDA) value (Figure 3A and Table S4). A total of 2,852 genes (cluster II in Figure 3A) were downregulated (>1.5-fold) in cells showing transcriptional repression and were not significantly regulated in other conditions. These genes are enriched in gene ontology pathways such as “cell cycle”, “RNA processing”, and “protein ubiquitination” (*P* < 10^−24^; Figure S4 and Figure 3A-B). The down-regulation of cell cycle genes indicates that the single-agent treatment inhibited cell proliferation. Especially, celecoxib treatment alone did not induce cell apoptosis, and the inhibition of cancer cell growth resulted from homogeneous inhibition of cell cycle genes across single cells (Figure 1B-C). Next, we examined whether the drug treatment modulated the expression of genes regulating a particular cell cycle phase. To this end, we analyzed signature genes uniquely expressed in the G1/S, S, G2, G2/M phases, respectively (24). These genesets showed a significant positive correlation of differential expression across single cells upon drug treatment (Figure S5). Although paclitaxel-treated cells are arrested in the G2/M phase (7), the inhibition of cell cycle genes is not limited to the particular phase.

**Figure 3.**
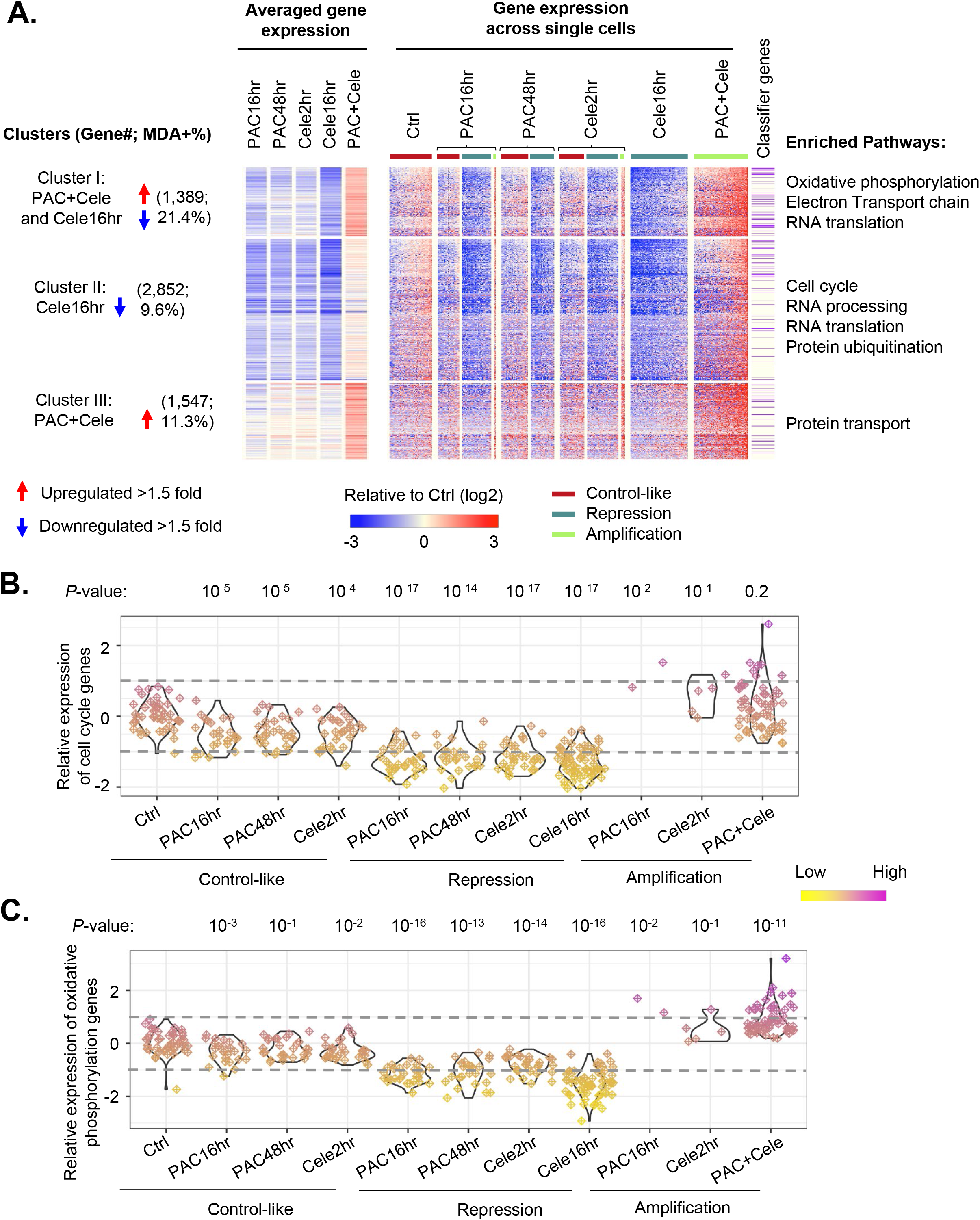
OXPHOS and cell cycle genes are major classifiers to define the cell states. (A) The heatmap showing the differentially expressed gene clusters based on their relative changes after drug treatment. The gene expression values were normalized to the mean of the control cells. Single cells in each treatment condition were further grouped by their defined states. The classifier genes in the random forest model with MDA value >0 are indicated in the heatmap. The enriched pathways in each cluster based on gene ontology analyses are shown. (B) The relative expression of cell cycle pathway genes across single cells, which were grouped based on drug treatment conditions and defined states. Each dot in the plot represents a single cell. The Wilcoxon rank sum test *P*-values comparing drug treatment groups vs. the control are shown. (C) Similar to B, the relative expression of OXPHOS pathway genes across single cells, which were grouped based on drug treatment conditions and defined states.

A total of 1,389 genes were down-regulated (>1.5-fold) in cells showing transcriptional repression by single drugs but were upregulated (>1.5-fold) upon combination treatment (cluster I in Figure 3A). These genes are enriched with MDA values >0 in the random forest model (Figure 3A) and contribute most significantly to cell state classification. Gene ontology analyses showed that they are enriched in pathways such as “OXPHOS”, “generation of precursor metabolites and energy”, and “RNA translation” (*P* < 10^−22^; Figure 3C and Figure S4). Unexpectedly, co-treatment of paclitaxel and celecoxib activated many genes regulating mitochondrial-related functions and the OXPHOS pathway. We further analyzed the genesets encoding complex I, II, III, IV, and ATP synthase of the electron transport chain. These genesets were synchronically regulated across single cells, suggesting a coherent transcriptional module regulates their expression during drug response (Figure S6). These results suggest that co-treatment with paclitaxel and celecoxib triggers a transcriptional amplification program activating OXPHOS, which can induce the generation of reactive oxygen species in cancer cells and promote cell apoptosis (31).

### Metabolism and inflammation genes are activated in a subpopulation of paclitaxel-treated cancer cells

At 48 h of paclitaxel treatment, 53% of cells showed global transcriptomic abundance comparable to untreated cells (Figure 2D). This subpopulation of cells did not fully mimic untreated cells, but showed unique gene signatures with 177 genes up-regulated by >1.5-fold (Figure 4A and Table S5). Among these, 20 genes show early activation (>1.5-fold) at 16 h, such as the interferon-induced transmembrane proteins IFITM1 and IFITM2, a driver of p53-dependent cell cycle arrest p21 (CDKN1A), and the growth differentiation factor-15 (GDF-15) (Figure 4B).

**Figure 4.**
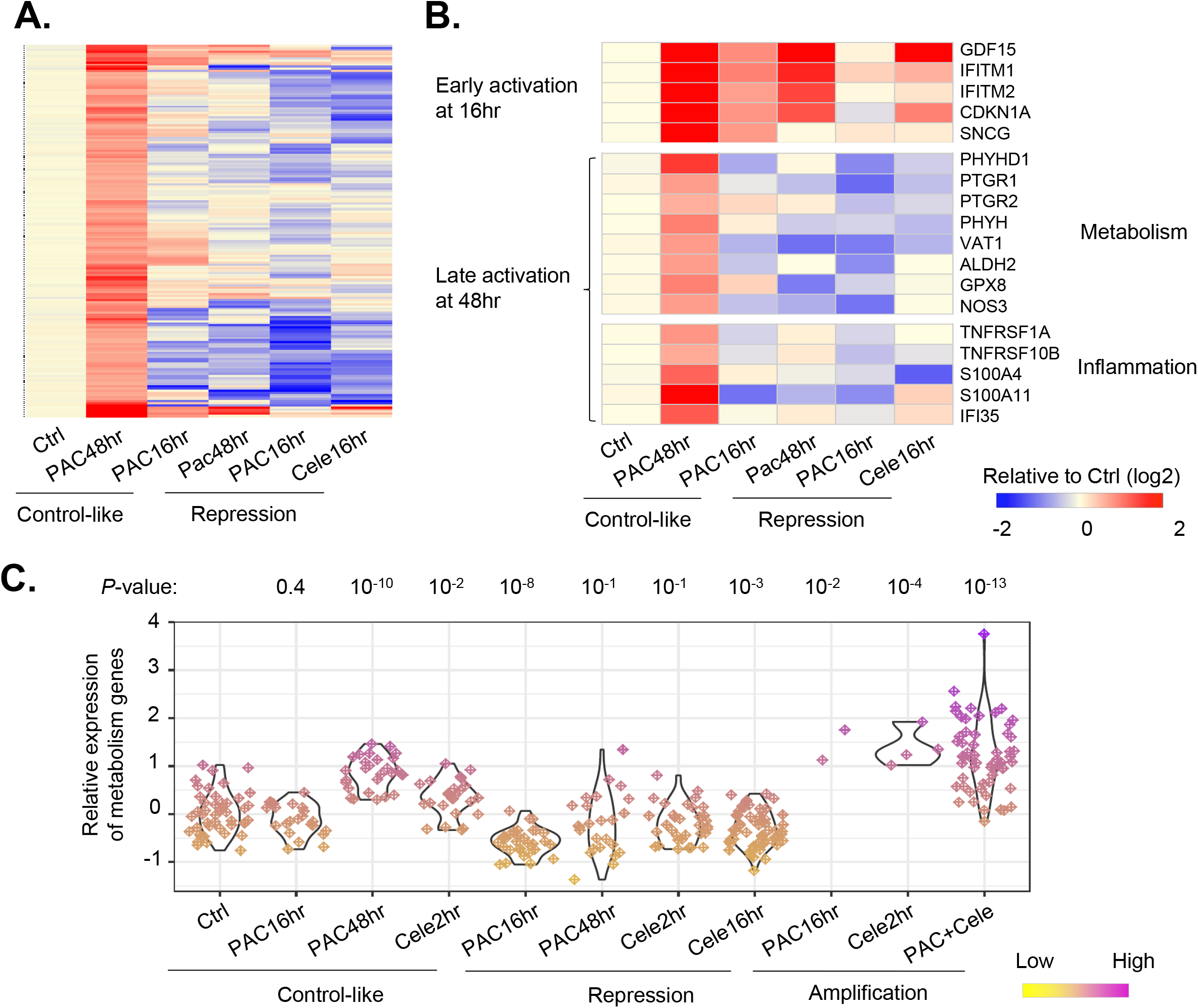
Genes activated in the control-like subpopulation of cells after 48 h of paclitaxel treatment. (A) Heatmap showing the expression changes of 177 genes activated in control-like cells after 48 h of paclitaxel treatment. Single cells were grouped based on drug treatment conditions and defined states. For each group, the averaged expression values of a gene across single cells were used to indicate its expression level. The expression fold changes compared to the control cells are shown in the heatmap. (B) Example paclitaxel-activated genes showing early activation at 16 h of paclitaxel treatment, and those showing late activation at 48 h and in pathways regulating metabolism and inflammation. (C) The relative expression of the metabolism genes across single cells grouped based on treatment conditions and defined states. The paclitaxel-activated genes in the GO:0055114: oxidation-reduction process were used for the analyses. The Wilcoxon rank sum test *P*-values comparing drug treatment groups vs. the control are shown.

The 157 other genes showed late activation at 48 h. The gene ontology analyses showed that they are enriched in genes regulating cellular metabolism (*P* < 10^−9^ for the pathway “oxidation-reduction (redox) process”) (Figure 4B and 4C). These include enzymes regulating lipid metabolism (e.g., ACACA, PTGR1, and SCD) and carboxylic acid biosynthesis (e.g. PHGDH and ASNS). In addition, some pro-inflammatory molecules, such as TNFRSF1A, IFI35, SERPINF1, IFNAR2, and S100A4, were up-regulated (Figure 4B). These results suggest that a subpopulation of cells rewired their intrinsic metabolic and inflammatory pathways after 48 h of paclitaxel treatment.

### A paclitaxel response index predicts treatment efficacy across several hundred cancer cell lines and in breast cancer patients

Next, we examined whether the activated inflammatory and metabolic gene signature plays a regulatory role in paclitaxel efficacy. We reasoned that if these genes function as a regulatory module, their co-activation should not be unique to paclitaxel response but should be general across biological conditions, because genes regulating a biological process tend to form a coexpression network (32,33). To this end, we examined the co-expression of the 177 paclitaxel-activated genes across 1,037 cancer cell lines from 26 primary tissue types using data from the Cancer Cell Line Encyclopedia (CCLE) database (25). By developing an iterative computational method, we found that a gene module consisting of 73 genes (41.2% of the total) showed a significant positive correlation of expression with each other (see Methods for details) (*P* < 10^−50^ compared to expected distribution; Figure 5A-B). The metabolic and inflammatory genes described above were present in this gene module. The data also indicate that the paclitaxel-induced gene module is intrinsically active in many cancer cells.

**Figure 5.**
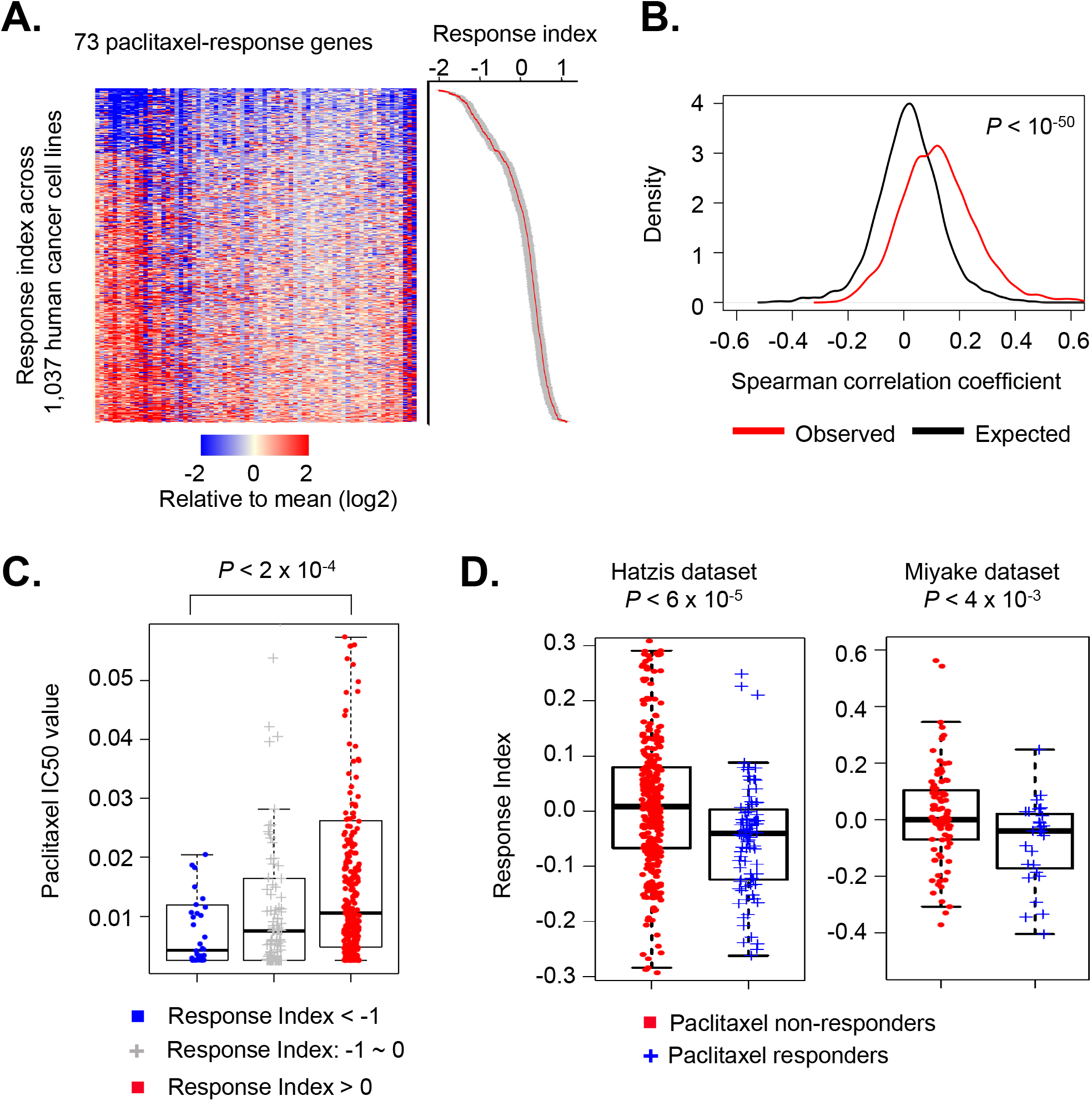
A gene module consisting of paclitaxel-activated genes predicts paclitaxel efficacy across cancer cell lines and breast cancer patients. (A) 73 paclitaxel-activated genes showed a positive correlation of expression across 1,037 cancer cell lines. We developed a paclitaxel response index to quantitatively measure the relative expression of these genes in a sample (standard error values are shown in grey). (B) Distribution of the observed and expected Spearman correlation coefficients of the expression levels between the gene pairs. The observed values were calculated based on gene pairs in (A). The expected correlation values were calculated by randomly picked gene pairs expressed in the cancer cell lines. The Wilcoxon rank sum test *P*-value comparing the observed vs. expected correlation coefficients is shown. (C) Cancer cell lines were grouped based on the paclitaxel-response index values in (A). Then we compared their paclitaxel IC_50_ values. The Wilcoxon Rank-Sum test *P*-value comparing two indicated groups is shown in the plot. (D) Breast cancer patients from two cohorts were grouped based on their clinical response to paclitaxel treatment. Then we compared the paclitaxel response index between the two groups of patients. The Wilcoxon Rank-Sum test *P*-values comparing two groups are shown in the plot.

As CCLE also measured the half-maximal inhibitory concentration (IC_50_) values of paclitaxel in these cancer cell lines, we examined whether there is a correlation between the gene module expression and the IC_50_ values. Based on the relative expression of the genes in the module, we developed a paclitaxel response index to quantify the geneset expression, which was calculated as the median of the normalized expression values of the 73 genes (Figure 5A). Next, we grouped cell lines based on their paclitaxel response index. Cancer cells with greater index values tended to show higher paclitaxel IC_50_ values (*P* < 2 × 10^−4^, Wilcoxon rank-sum test; Figure 5C and Table S6). These data indicate that higher baseline expression of the gene module is linked with decreased paclitaxel efficacy in cancer cells.

Furthermore, we examined the impact of the gene module on paclitaxel response across breast cancer patients. We analyzed the RNA expression and clinical data from two published breast cancer patient cohorts (26,27). Indeed, responders to paclitaxel treatment showed lower paclitaxel response index values compared to non-responders (Figures 5D and S7, Table S7). These results further indicate that our paclitaxel response index can predict the clinical outcome of paclitaxel treatment.

## DISCUSSION

Using the Fluidigm C1 Single-Cell Auto Prep System, we added a constant amount of the spike-in RNA set to each well during library preparation. During the data analyses, we calculated gene expression levels by aligning sequencing reads to a hybrid transcriptome combining genome-encoded transcripts and spike-in RNAs. The fold differences of spike-in RNAs learned from the sequencing data are in accordance with the pre-designed concentrations, indicating that our scRNA-seq quantitatively measures RNA levels. Furthermore, by normalizing endogenous gene expression to that of spike-in RNAs, we found that mRNA abundance is drastically differentially regulated across single cells after drug treatment. These differences would not be observed in measurements using TPM or RPKM (reads per kilobase of transcript per million reads mapped) values for endogenous genes alone, as these calculations rely on the presumption that global transcriptomic levels are comparable across cells. Our results confirm the value of adding the synthetic spike-in RNAs for scRNA-seq.

One of the major findings of this study is that drug treatments induce variable changes in global mRNA abundance in a cell. This level of regulation has been commonly overlooked by previous studies due to the lack of gene expression normalization using spike-in RNAs for RNA-seq experiments. Interestingly, paclitaxel treatment alone induces transcriptional repression in a subpopulation of cells. The down-regulation of cell cycle genes is consistent with the mitotic arrest induced by paclitaxel. Here we found that genes expressed in the G1/S, S, G2, G2/M phases are synchronically downregulated at comparable levels, indicating the inhibition of cell cycle genes is not specific to a particular phase. A coordinated transcriptional network is likely to regulate the expression of cell cycle genes.

Interestingly, the combination of paclitaxel and celecoxib induces transcriptional amplification, which is opposite to the transcriptional repression caused by the single drug treatments (Figure 6). The fact that the expression of cell cycle genes is unchanged after co-treatment with the two drugs suggests that another regulatory pathway promotes cell apoptosis in this context. Unexpectedly, the OXPHOS pathway showed the most significant up-regulation. Hyperactivation of OXPHOS can cause leakage of electrons from electron transport chains, leading to a partial reduction of oxygen and the formation of superoxide in cells (34). This could be the mechanism by which the drug combination enhances the cell-killing efficiency. These results also indicate that the interplay between two drugs can trigger a novel transcriptional program, increasing the efficacy.

**Figure 6.**
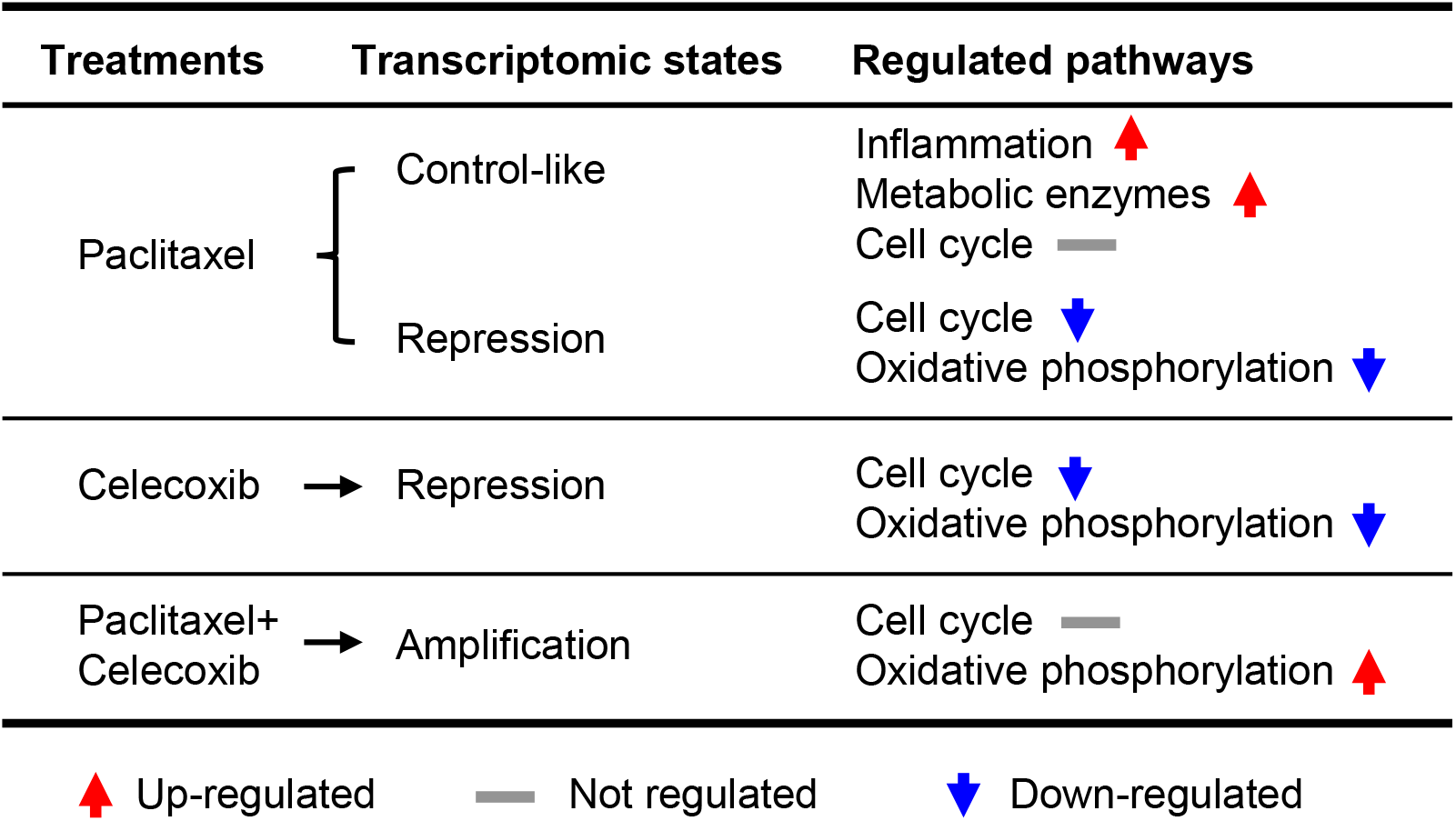
The table shows the transcriptomic states and regulation of associated pathways after the various drug treatments.

Dynamic global transcriptional activity can result from the regulation of the cellular amount of RNA polymerase II (Pol II), the efficiency of Pol II recruitment to promoters, or the rate of transcriptional elongation (35,36). These molecular mechanisms have been characterized in a few biological processes. Cancer cells with overexpressed MYC show higher transcriptomic abundance than normal cells because MYC, located in promoters and enhancers can directly recruit the transcriptional elongation factor P-TEFb, resulting in transcriptional amplification (37). In B-cell acute lymphoblastic leukemia, overexpression of the nucleosome remodeling protein HMGN1 suppresses heterochromatin marker H3K27me3 levels and promotes B cell transcriptomic levels and proliferation (38). Future efforts are needed to identify the molecular regulators mediating transcriptional amplification or suppression after drug treatment.

The initial expression changes of cancer cells in response to drugs are important in determining their fate. The probability of drug-induced apoptosis appears to be stochastic from the perspective of a single cell, but has a fixed ratio in the larger cell population (39–41). The preexisting gene expression programs regulate the ratio of killed cells (i.e. the IC_50_ value of a drug). At the single-cell level, cell death is determined by whether pro-apoptotic signals have accumulated to the threshold level. Here we found that transcriptomic abundance shows intercellular heterogeneity in response to paclitaxel treatment. This heterogeneity is a new regulatory layer contributing to chemoresistance. In about half of the treated cells, the global transcriptional levels are comparable to the control state. Unexpectedly, a coherent gene module consisting of a small number of genes (73 genes) was activated in a subpopulation of paclitaxel-treated cells, and these genes were highly enriched in redox and inflammatory pathways. We showed that the baseline expression of this gene module predicts chemo-response *in vitro* and *in vivo*. The metabolic switch in cancer cells is known to play a major regulatory role during drug response (42,43). For example, some genes in the module, such as aldehyde dehydrogenase (ALDH2) and acetyl-CoA carboxylase (ACACA), are known oncogenes, promoting cancer stem cell formation and drug resistance (44,45). Future experiments tracing the metabolic changes of cancer cells after drug treatment will provide further functional insights into this level of regulation.

## ACKNOWLEDGMENTS

This work was supported by the following grants to ZJ: the Searle Leadership Fund in the Life Sciences from Northwestern University, the National Cancer Institute (R00CA207865), the Lynn Sage Scholar grant from the Lynn Sage Foundation, and the Lynn Sage Cancer Research Foundation. VB is supported by the National Insitutes of Health (R33CA225323 and R01CA225002) and the National Science Foundation (EFMA-1830961). We thank Marcus Peter, Daniela Matei, and the members of the Ji lab for helpful discussions.

## AVAILABILITY OF DATA AND MATERIALS

The sequencing datasets generated during the current study are available in the Gene Expression Omnibus (GEO) repository with the accession number GSE162256. The following secure token has been created to allow review of record GSE162256 while it remains in private status: mbcbsqgczpatjch. Computational codes are available upon request.

## REFERENCES

1. Ben-David, U., Siranosian, B., Ha, G., Tang, H., Oren, Y., Hinohara, K., Strathdee, C.A., Dempster, J., Lyons, N.J., Burns, R. et al. (2018) Genetic and transcriptional evolution alters cancer cell line drug response. Nature, 560, 325–330.

2. Holohan, C., Van Schaeybroeck, S., Longley, D.B. and Johnston, P.G. (2013) Cancer drug resistance: an evolving paradigm. Nat Rev Cancer, 13, 714–726.

3. Rathert, P., Roth, M., Neumann, T., Muerdter, F., Roe, J.S., Muhar, M., Deswal, S., Cerny-Reiterer, S., Peter, B., Jude, J. et al. (2015) Transcriptional plasticity promotes primary and acquired resistance to BET inhibition. Nature, 525, 543–547.

4. Flavahan, W.A., Gaskell, E. and Bernstein, B.E. (2017) Epigenetic plasticity and the hallmarks of cancer. Science, 357.

5. Jiang, P., Sellers, W.R. and Liu, X.S. (2018) Big Data Approaches for Modeling Response and Resistance to Cancer Drugs. Annu Rev Biomed Data Sci, 1, 1–27.

6. Weaver, B.A. (2014) How Taxol/paclitaxel kills cancer cells. Mol Biol Cell, 25, 2677–2681.

7. Yusuf, R.Z., Duan, Z., Lamendola, D.E., Penson, R.T. and Seiden, M.V. (2003) Paclitaxel resistance: molecular mechanisms and pharmacologic manipulation. Curr Cancer Drug Targets, 3, 1–19.

8. Orr, G.A., Verdier-Pinard, P., McDaid, H. and Horwitz, S.B. (2003) Mechanisms of Taxol resistance related to microtubules. Oncogene, 22, 7280–7295.

9. Yang, H., Mao, W., Rodriguez-Aguayo, C., Mangala, L.S., Bartholomeusz, G., Iles, L.R., Jennings, N.B., Ahmed, A.A., Sood, A.K., Lopez-Berestein, G. et al. (2018) Paclitaxel Sensitivity of Ovarian Cancer Can be Enhanced by Knocking Down Pairs of Kinases that Regulate MAP4 Phosphorylation and Microtubule Stability. Clin Cancer Res, 24, 5072–5084.

10. Vuylsteke, P., Huizing, M., Petrakova, K., Roylance, R., Laing, R., Chan, S., Abell, F., Gendreau, S., Rooney, I., Apt, D. et al. (2016) Pictilisib PI3Kinase inhibitor (a phosphatidylinositol 3-kinase [PI3K] inhibitor) plus paclitaxel for the treatment of hormone receptor-positive, HER2-negative, locally recurrent, or metastatic breast cancer: interim analysis of the multicentre, placebo-controlled, phase II randomised PEGGY study. Ann Oncol, 27, 2059–2066.

11. Kim, H.J., Yim, G.W., Nam, E.J. and Kim, Y.T. (2014) Synergistic Effect of COX-2 Inhibitor on Paclitaxel-Induced Apoptosis in the Human Ovarian Cancer Cell Line OVCAR-3. Cancer Res Treat, 46, 81–92.

12. Altorki, N.K., Keresztes, R.S., Port, J.L., Libby, D.M., Korst, R.J., Flieder, D.B., Ferrara, C.A., Yankelevitz, D.F., Subbaramaiah, K., Pasmantier, M.W. et al. (2003) Celecoxib, a selective cyclo-oxygenase-2 inhibitor, enhances the response to preoperative paclitaxel and carboplatin in early-stage non-small-cell lung cancer. J Clin Oncol, 21, 2645–2650.

13. Olsen, S.R. (2005) Taxanes and COX-2 inhibitors: from molecular pathways to clinical practice. Biomed Pharmacother, 59 Suppl 2, S306–310.

14. Almassalha, L.M., Bauer, G.M., Wu, W., Cherkezyan, L., Zhang, D., Kendra, A., Gladstein, S., Chandler, J.E., VanDerway, D., Seagle, B.L. et al. (2017) Macrogenomic engineering via modulation of the scaling of chromatin packing density. Nat Biomed Eng, 1, 902–913.

15. Picelli, S., Bjorklund, A.K., Faridani, O.R., Sagasser, S., Winberg, G. and Sandberg, R. (2013) Smart-seq2 for sensitive full-length transcriptome profiling in single cells. Nat Methods, 10, 1096–1098.

16. Lun, A.T.L., Calero-Nieto, F.J., Haim-Vilmovsky, L., Gottgens, B. and Marioni, J.C. (2017) Assessing the reliability of spike-in normalization for analyses of single-cell RNA sequencing data. Genome Res, 27, 1795–1806.

17. Li, B. and Dewey, C.N. (2011) RSEM: accurate transcript quantification from RNA-Seq data with or without a reference genome. BMC Bioinformatics, 12, 323.

18. Frankish, A., Diekhans, M., Ferreira, A.M., Johnson, R., Jungreis, I., Loveland, J., Mudge, J.M., Sisu, C., Wright, J., Armstrong, J. et al. (2019) GENCODE reference annotation for the human and mouse genomes. Nucleic Acids Res, 47, D766–D773.

19. Langmead, B. and Salzberg, S.L. (2012) Fast gapped-read alignment with Bowtie 2. Nat Methods, 9, 357–359.

20. Puram, S.V., Tirosh, I., Parikh, A.S., Patel, A.P., Yizhak, K., Gillespie, S., Rodman, C., Luo, C.L., Mroz, E.A., Emerick, K.S. et al. (2017) Single-Cell Transcriptomic Analysis of Primary and Metastatic Tumor Ecosystems in Head and Neck Cancer. Cell, 171, 1611–1624 e1624.

21. Tirosh, I., Izar, B., Prakadan, S.M., Wadsworth, M.H., 2nd, Treacy, D., Trombetta, J.J., Rotem, A., Rodman, C., Lian, C., Murphy, G. et al. (2016) Dissecting the multicellular ecosystem of metastatic melanoma by single-cell RNA-seq. Science, 352, 189–196.

22. Eisenberg, E. and Levanon, E.Y. (2013) Human housekeeping genes, revisited. Trends Genet, 29, 569–574.

23. Huang da, W., Sherman, B.T. and Lempicki, R.A. (2009) Systematic and integrative analysis of large gene lists using DAVID bioinformatics resources. Nat Protoc, 4, 44–57.

24. Whitfield, M.L., Sherlock, G., Saldanha, A.J., Murray, J.I., Ball, C.A., Alexander, K.E., Matese, J.C., Perou, C.M., Hurt, M.M., Brown, P.O. et al. (2002) Identification of genes periodically expressed in the human cell cycle and their expression in tumors. Mol Biol Cell, 13, 1977–2000.

25. Ghandi, M., Huang, F.W., Jane-Valbuena, J., Kryukov, G.V., Lo, C.C., McDonald, E.R., 3rd, Barretina, J., Gelfand, E.T., Bielski, C.M., Li, H. et al. (2019) Next-generation characterization of the Cancer Cell Line Encyclopedia. Nature, 569, 503–508.

26. Hatzis, C., Pusztai, L., Valero, V., Booser, D.J., Esserman, L., Lluch, A., Vidaurre, T., Holmes, F., Souchon, E., Wang, H. et al. (2011) A genomic predictor of response and survival following taxane-anthracycline chemotherapy for invasive breast cancer. JAMA, 305, 1873–1881.

27. Miyake, T., Nakayama, T., Naoi, Y., Yamamoto, N., Otani, Y., Kim, S.J., Shimazu, K., Shimomura, A., Maruyama, N., Tamaki, Y. et al. (2012) GSTP1 expression predicts poor pathological complete response to neoadjuvant chemotherapy in ER-negative breast cancer. Cancer Sci, 103, 913–920.

28. Ziegenhain, C., Vieth, B., Parekh, S., Reinius, B., Guillaumet-Adkins, A., Smets, M., Leonhardt, H., Heyn, H., Hellmann, I. and Enard, W. (2017) Comparative Analysis of Single-Cell RNA Sequencing Methods. Mol Cell, 65, 631–643 e634.

29. Paek, A.L., Liu, J.C., Loewer, A., Forrester, W.C. and Lahav, G. (2016) Cell-to-Cell Variation in p53 Dynamics Leads to Fractional Killing. Cell, 165, 631–642.

30. Vallejos, C.A., Risso, D., Scialdone, A., Dudoit, S. and Marioni, J.C. (2017) Normalizing single-cell RNA sequencing data: challenges and opportunities. Nat Methods, 14, 565–571.

31. Ashton, T.M., McKenna, W.G., Kunz-Schughart, L.A. and Higgins, G.S. (2018) Oxidative Phosphorylation as an Emerging Target in Cancer Therapy. Clin Cancer Res, 24, 2482–2490.

32. Stuart, J.M., Segal, E., Koller, D. and Kim, S.K. (2003) A gene-coexpression network for global discovery of conserved genetic modules. Science, 302, 249–255.

33. Ji, Z., He, L., Regev, A. and Struhl, K. (2019) Inflammatory regulatory network mediated by the joint action of NF-kB, STAT3, and AP-1 factors is involved in many human cancers. Proc Natl Acad Sci U S A, 116, 9453–9462.

34. Li, X., Fang, P., Mai, J., Choi, E.T., Wang, H. and Yang, X.F. (2013) Targeting mitochondrial reactive oxygen species as novel therapy for inflammatory diseases and cancers. J Hematol Oncol, 6, 19.

35. Li, Y., Wang, H., Muffat, J., Cheng, A.W., Orlando, D.A., Loven, J., Kwok, S.M., Feldman, D.A., Bateup, H.S., Gao, Q. et al. (2013) Global transcriptional and translational repression in human-embryonic-stem-cell-derived Rett syndrome neurons. Cell Stem Cell, 13, 446–458.

36. Loven, J., Orlando, D.A., Sigova, A.A., Lin, C.Y., Rahl, P.B., Burge, C.B., Levens, D.L., Lee, T.I. and Young, R.A. (2012) Revisiting global gene expression analysis. Cell, 151, 476–482.

37. Lin, C.Y., Loven, J., Rahl, P.B., Paranal, R.M., Burge, C.B., Bradner, J.E., Lee, T.I. and Young, R.A. (2012) Transcriptional amplification in tumor cells with elevated c-Myc. Cell, 151, 56–67.

38. Lane, A.A., Chapuy, B., Lin, C.Y., Tivey, T., Li, H., Townsend, E.C., van Bodegom, D., Day, T.A., Wu, S.C., Liu, H. et al. (2014) Triplication of a 21q22 region contributes to B cell transformation through HMGN1 overexpression and loss of histone H3 Lys27 trimethylation. Nat Genet, 46, 618–623.

39. Roux, J., Hafner, M., Bandara, S., Sims, J.J., Hudson, H., Chai, D. and Sorger, P.K. (2015) Fractional killing arises from cell-to-cell variability in overcoming a caspase activity threshold. Mol Syst Biol, 11, 803.

40. Cohen, A.A., Geva-Zatorsky, N., Eden, E., Frenkel-Morgenstern, M., Issaeva, I., Sigal, A., Milo, R., Cohen-Saidon, C., Liron, Y., Kam, Z. et al. (2008) Dynamic proteomics of individual cancer cells in response to a drug. Science, 322, 1511–1516.

41. Spencer, S.L., Gaudet, S., Albeck, J.G., Burke, J.M. and Sorger, P.K. (2009) Non-genetic origins of cell-to-cell variability in TRAIL-induced apoptosis. Nature, 459, 428–432.

42. Ratnikov, B.I., Scott, D.A., Osterman, A.L., Smith, J.W. and Ronai, Z.A. (2017) Metabolic rewiring in melanoma. Oncogene, 36, 147–157.

43. Rahman, M. and Hasan, M.R. (2015) Cancer Metabolism and Drug Resistance. Metabolites, 5, 571–600.

44. Raha, D., Wilson, T.R., Peng, J., Peterson, D., Yue, P., Evangelista, M., Wilson, C., Merchant, M. and Settleman, J. (2014) The cancer stem cell marker aldehyde dehydrogenase is required to maintain a drug-tolerant tumor cell subpopulation. Cancer Res, 74, 3579–3590.

45. Chajes, V., Cambot, M., Moreau, K., Lenoir, G.M. and Joulin, V. (2006) Acetyl-CoA carboxylase alpha is essential to breast cancer cell survival. Cancer Res, 66, 5287–5294.

