## Supplemental Figures for "Spike-in normalization for single-cell RNA-seq reveals dynamic global transcriptional activity mediating anti-cancer drug response"

#### SUPPLEMENTARY INFORMATION

**Figure S1. Quality control of scRNA-seq.**

**Figure S2. The relative expression of 3 spike-in molecules across single cells.**

**Figure S3. The expression ratio between endogenous genes and the spike-in molecule.**

**Figure S4. The gene ontology analyses of 3 gene clusters in Figure 3A.**

**Figure S5. The relative expression of cell cycle genes across single cells.**

**Figure S6. The relative expression of 5 complexes of electron transport chain genes across single cells.**

**Figure S7. The Paclitaxel-response index across breast cancer patients.**

**Table S1. Sequences of the three spike-in RNAs.**

**Table S2. The statistics of scRNA-seq experiments and analyses.**

**Table S3. Three clusters of genes showing significant expression regulation after drug treatment as in Figure 3A.**

**Table S4. Classifier genes with MDA values greater than 0 from the random forest model.**

**Table S5. 177 genes activated in the control-like subpopulation of paclitaxel-treated cells (48 h).**

**Table S6. Paclitaxel response index across 1,037 human cancer cell lines.**

**Table S7. Paclitaxel response index across human patient samples, including Hatzis and Miyake datasets.**

### Figure S1

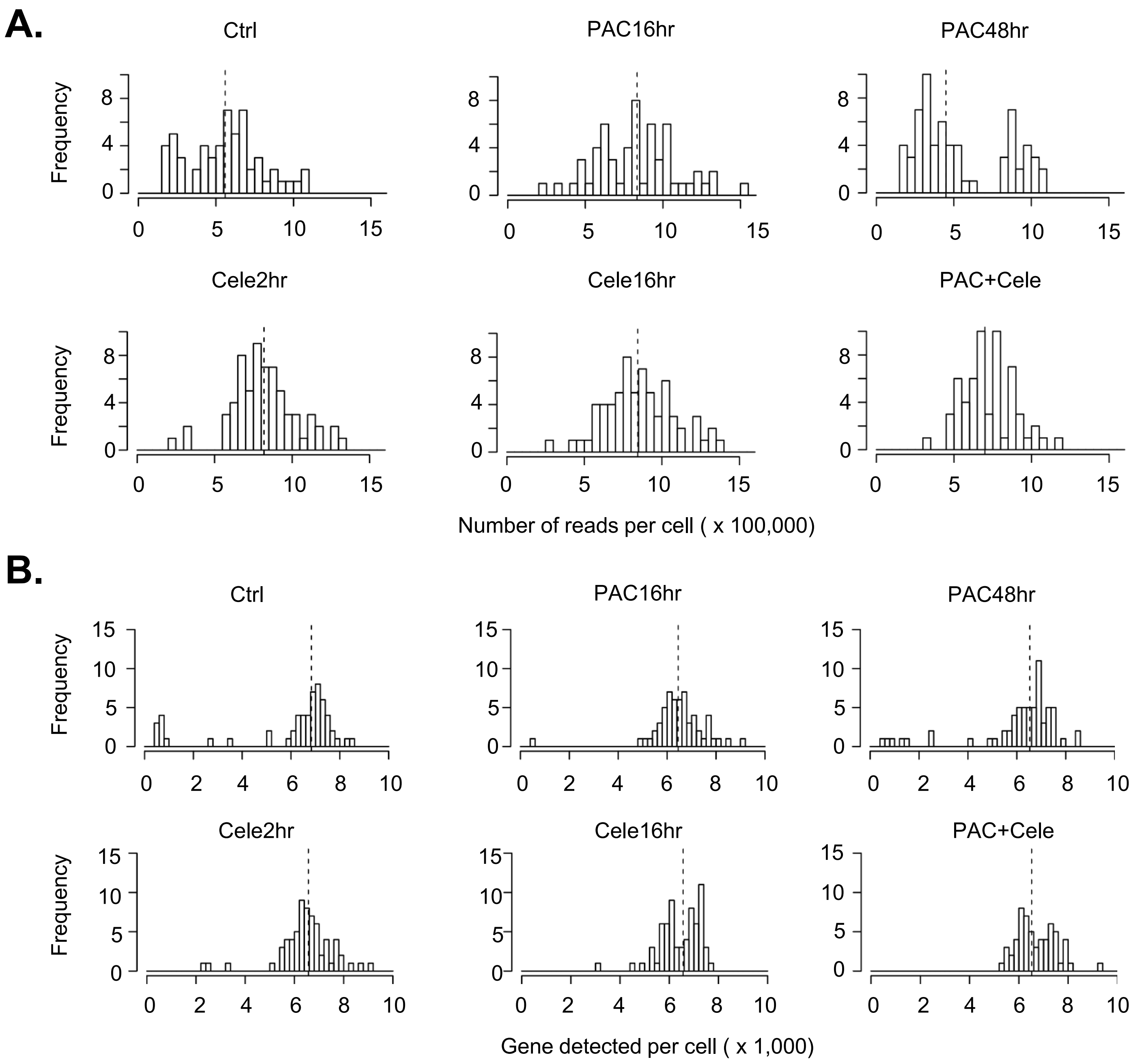


**Figure S1. Quality control of scRNA-seq.**
(A) The distribution of the read number per cell.
(B) The distribution of the number of genes detected per cell.

### Figure S2

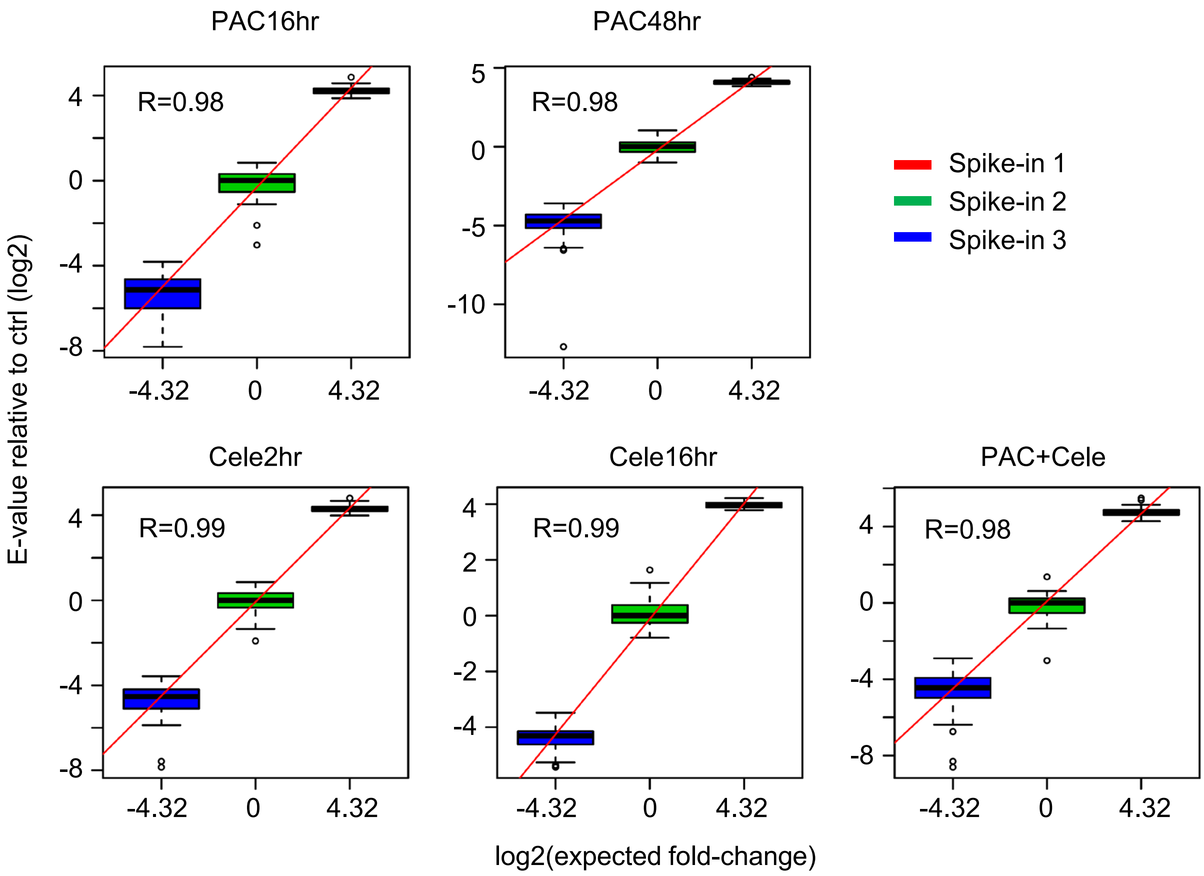


**Figure S2. The relative expression of 3 spike-in molecules across single cells.** The X-axis indicates the expected expression differences and Y-axis shows fold differences measured by scRNA-seq. The values were normalized to the median of the spike-in 2 expression. The Pearson correlation coefficients are indicated in the plots.

### Figure S3

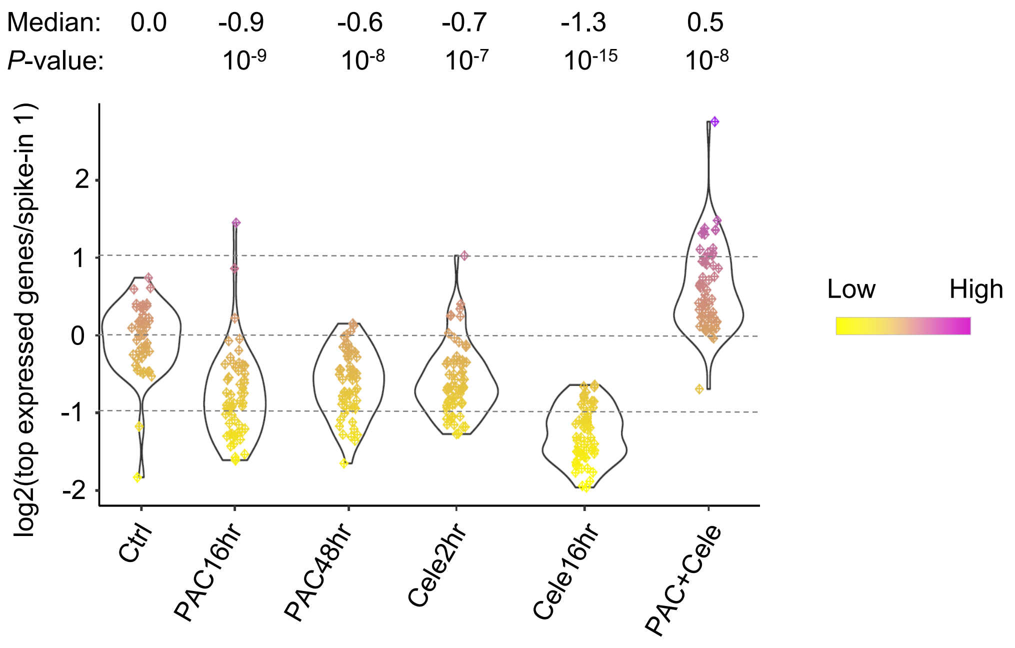


**Figure S3. The expression ratio between endogenous genes and the spike-in molecule.** We used the top 5,000 expressed genes in a single cell to measure the overall endogenous gene expression and used the highest expressed spike-in molecule (spike-in 1) to indicate spike-in RNA expression. The log2(ratio) values were then normalized to the median of the control cells.

### Figure S4

| | GO term | Cluster I | Cluster II | Cluster III |
|---|---|---|---|---|
| Biological process | GO:0006412 translation | 35.2 | 36.6 | 0.1 |
| | GO:0006414 translational elongation | 28.6 | 9.3 | 0.0 |
| | GO:0006091 generation of precursor metabolites and energy | 27.6 | 5.4 | 0.2 |
| | GO:0045333 cellular respiration | 24.2 | 2.0 | 0.1 |
| | GO:0006119 oxidative phosphorylation | 22.9 | 2.6 | 0.0 |
| | GO:0006396 RNA processing | 4.8 | 75.3 | 0.0 |
| | GO:0007049 cell cycle | 0.8 | 56.0 | 0.4 |
| | GO:0048285 organelle fission | 1.3 | 36.2 | 0.0 |
| | GO:0051340 regulation of ligase activity | 0.9 | 29.4 | 0.0 |
| | GO:0051438 regulation of ubiquitin-protein ligase activity | 0.9 | 27.4 | 0.0 |
| | GO:0051301 cell division | 1.2 | 26.2 | 0.1 |
| | GO:0031396 regulation of protein ubiquitination | 1.2 | 24.1 | 0.1 |
| | GO:0008104 protein localization | 11.4 | 7.7 | 6.4 |
| | GO:0015031 protein transport | 12.2 | 8.0 | 5.7 |
| Cellular component | GO:0005739 mitochondrion | 56.3 | 42.2 | 1.5 |
| | GO:0005840 ribosome | 32.2 | 29.6 | 0.0 |
| | GO:0031090 organelle membrane | 29.1 | 4.7 | 5.3 |
| | GO:0030529 ribonucleoprotein complex | 28.9 | 83.8 | 0.0 |
| | GO:0070013 intracellular organelle lumen | 12.2 | 121.3 | 1.3 |
| | GO:0005654 nucleoplasm | 2.9 | 73.1 | 0.1 |
| | GO:0005681 spliceosome | 2.4 | 34.6 | 0.0 |
| | GO:0005694 chromosome | 0.7 | 31.1 | 0.0 |
| | GO:0000502 proteasome complex | 0.5 | 23.4 | 0.0 |
| | GO:0005794 Golgi apparatus | 3.6 | 0.0 | 13.4 |
| | GO:0005783 endoplasmic reticulum | 10.3 | 0.0 | 9.9 |
| Molecular function | GO:0003735 structural constituent of ribosome | 31.9 | 23.3 | 0.0 |
| | GO:0003954 NADH dehydrogenase activity | 18.8 | 0.8 | 0.0 |
| | GO:0016651 oxidoreductase activity, acting on NADH or NADPH | 18.3 | 0.8 | 0.1 |
| | GO:0003723 RNA binding | 15.3 | 52.5 | 0.1 |
| | GO:0000166 nucleotide binding | 3.3 | 20.7 | 2.3 |
| | GO:0005524 ATP binding | 0.1 | 13.9 | 3.6 |
| | GO:0003899 DNA-directed RNA polymerase activity | 0.1 | 11.3 | 0.0 |
| | GO:0034062 RNA polymerase activity | 0.1 | 11.3 | 0.0 |
| | GO:0016779 nucleotidyltransferase activity | 0.4 | 11.0 | 0.6 |
| | GO:0016887 ATPase activity | 0.9 | 10.9 | 1.9 |
| | GO:0016879 ligase activity, forming carbon-nitrogen bonds | 1.6 | 3.9 | 4.3 |

-log10(*P*-value)

0 — 30

**Figure S4. The gene ontology analyses of the 3 gene clusters in Figure 3A.** The –log10(P-values) of each pathway in the 3 groups are shown.

### Figure S5

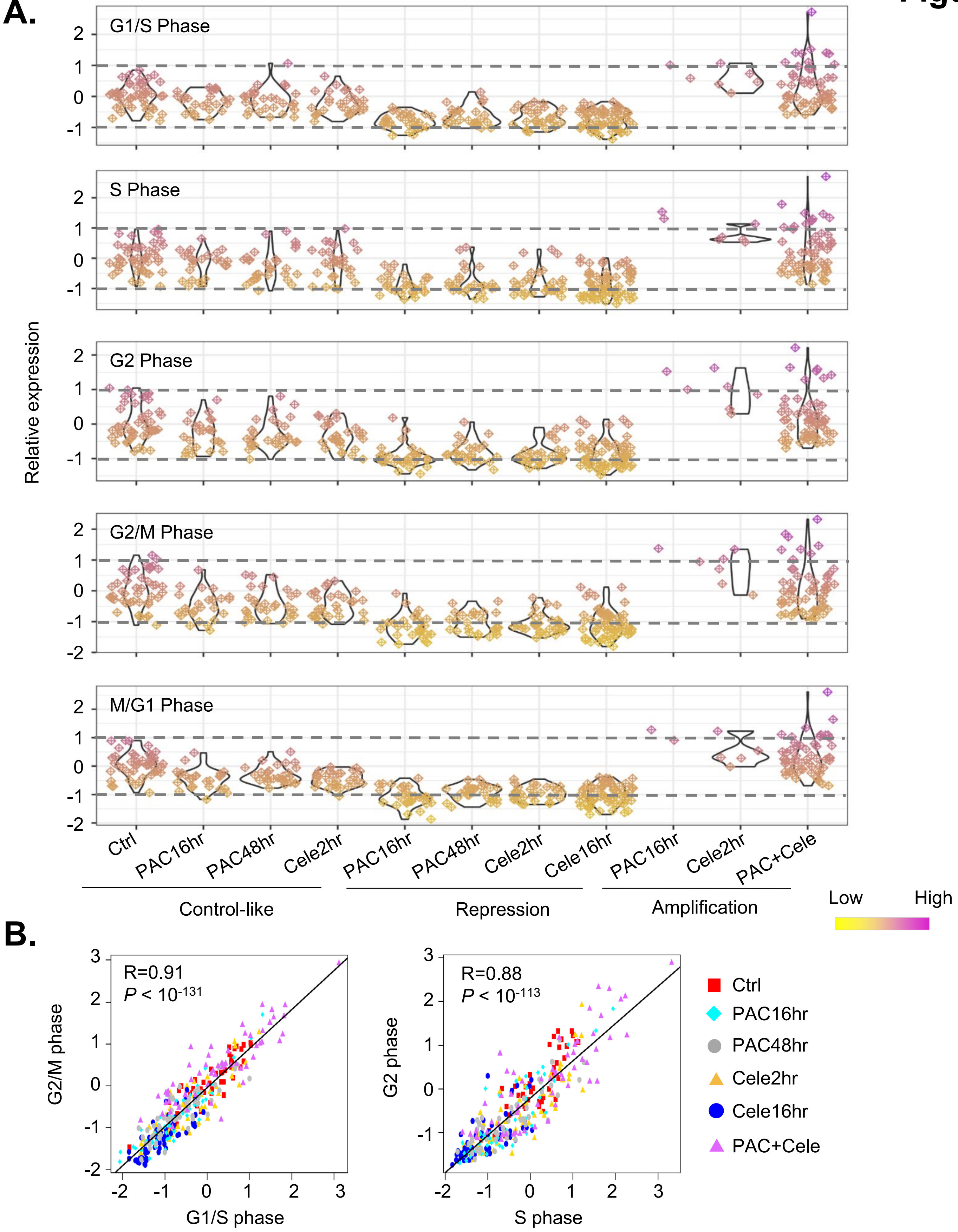


**Figure S5. The relative expression of cell cycle genes across single cells.**

(A) The relative expression of genesets regulating G1/S phase, S phase, G2 phase, G2/M phase, and M/G1 phase. The single cells were grouped based on drug treatment conditions and defined states.

(B) The correlation between differential expression levels of genesets regulating different cell cycle phases. The Pearson correlation coefficient values and the linear regression *P*-values are shown.

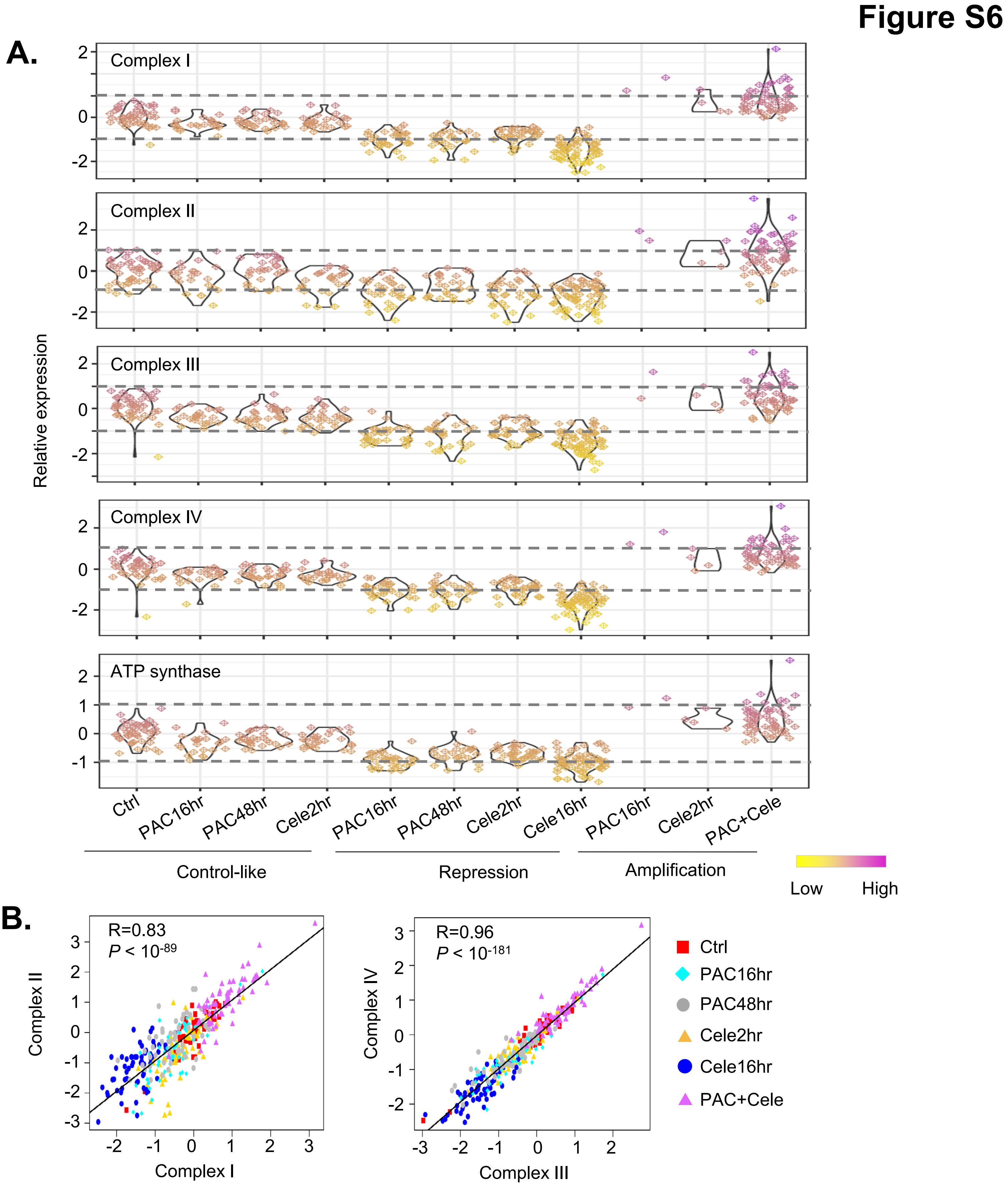


**Figure S6. The relative expression of 5 electron transport chain complex genes across single cells.**

(A) The relative expression of genesets encoding complex I, complex II, complex III, complex IV, and complex V (ATP synthase). The single cells were grouped based on drug treatment conditions and defined states.

(B) The correlation between differential expression levels of genesets encoding the complexes. The Pearson correlation coefficient values and the linear regression *P*-values are shown.

### Figure S7

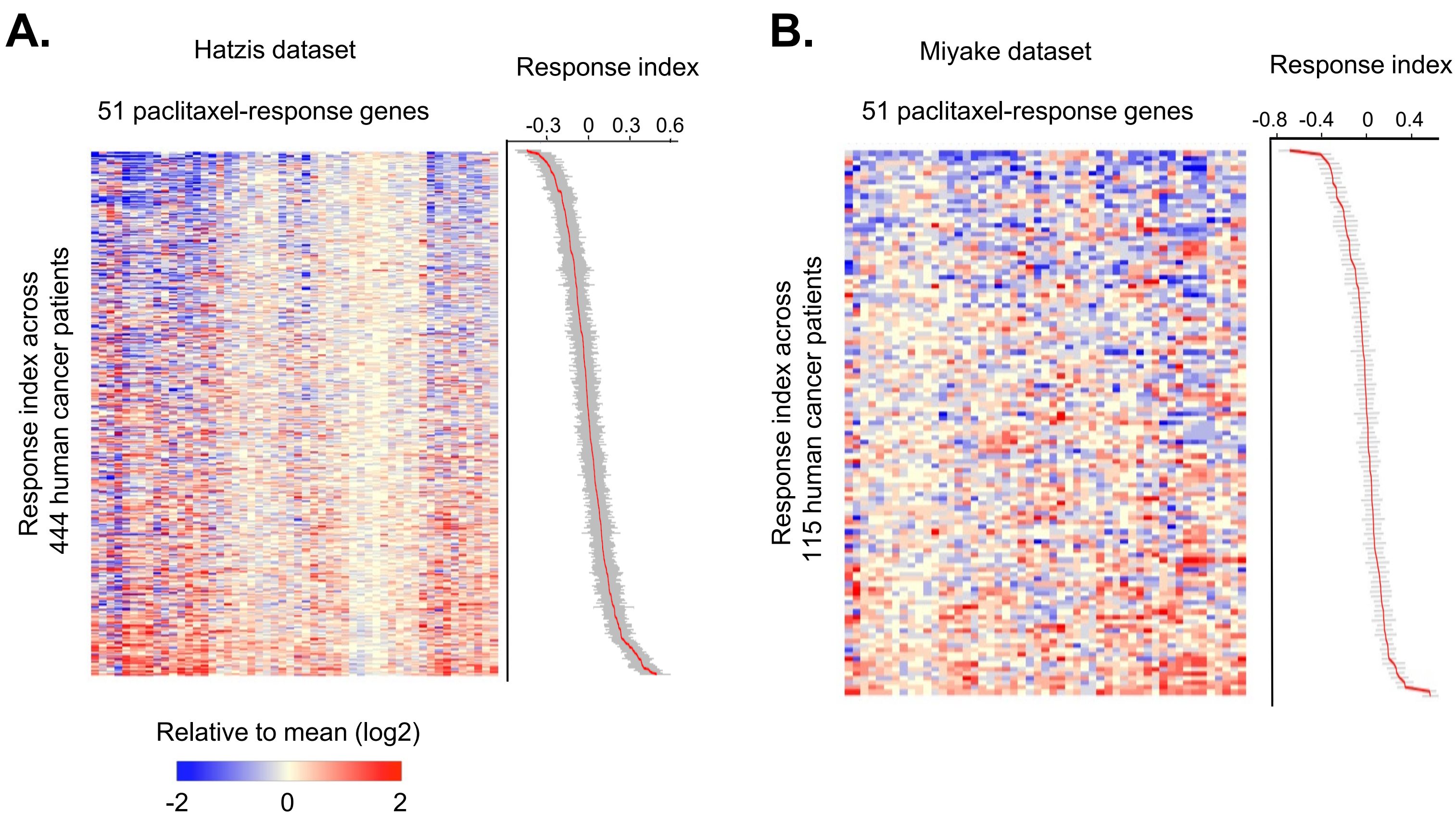


**Figure S7. The paclitaxel response index across breast cancer patients.** The relative expression of 51 paclitaxel-activated genes across breast cancer patients' samples using the Hatzis dataset (A) and the Miyake dataset (B). We developed a paclitaxel response index to quantitatively measure the relative expression of these genes in a sample (standard error values are shown in grey).
